# Informing generative replay for continual learning with long-term memory formation in the fruit fly

**DOI:** 10.1101/2023.01.18.524467

**Authors:** Brian S. Robinson, Justin Joyce, Raphael Norman-Tenazas, Gautam K. Vallabha, Erik C. Johnson

## Abstract

Continual learning without catastrophic forgetting is a challenge for artificial systems but it is done naturally across a range of biological systems, including in insects. A recurrent circuit has been identified in the fruit fly mushroom body to consolidate long term memories (LTM), but there is not currently an algorithmic understanding of this LTM formation. We hypothesize that generative replay is occurring to consolidate memories in this recurrent circuit, and find anatomical evidence in synapse-level connectivity that supports this hypothesis. Next, we introduce a computational model which combines an initial experience phase and LTM phase to perform generative replay based continual learning. When evaluated on a CIFAR-100 class-incremental continual learning task, the modeled LTM phase increases classification performance by 20% and approaches within 2% of the performance for a non-incremental upper bound baseline. Unique elements of the proposed generative replay model include: 1) coupling high dimensional sparse activation patterns with generative replay and 2) sampling and reconstructing higher level representations for training generative replay (as opposed to reconstructing sensory-level or processed sensory-level representations). Additionally, we make the experimentally testable prediction that a specific set of synapses would need to undergo experience-dependent plasticity during LTM formation to support our generative replay based model.

## 1 Introduction

Despite the high performance of deep learning systems for classification tasks, continual learning without catastrophic forgetting of prior classes remains a challenge for deep learning systems. Current approaches to address this [10] include structural, regularization, and replay-based, with replay-based being the most effective at combating catastrophic forgetting and broadly attributed to biological systems [3]. A challenge with existing replay-based continual learning approaches, however, is the increasing memory and compute time required as number of tasks learned increases. Another challenge is learning from a single experience of data, as buffer based replay and generative replay networks require several epochs of training with recently stored data. While observations and system level principles have been identified for how mammals perform replay for lifelong learning, it is difficult to study the underlying neural circuit mechanisms with synapse-level detail, which can pose limitations to informing artificial continual learning systems.

A model system for biological learning is the fruit fly, where the mushroom body (the learning center) is well characterized [4; 9] and unprecedented detailed synapse-level connectivity has been recently made available [11; 7]. The feedforward path of this system is the most understood [4], in which processed sensory input encoded in projection neuron (PN) activity is projected to a higher dimensional sparse representation in Kenyon Cells (KC’s) which subsequently synapse with a smaller population of mushroom body output neurons (MBON’s). MBON activation drives higher-level behavioral decisions and associative learning at the synapses between KC’s and MBON’s is a substrate for learning to link sensory input patterns with rewarded behavioral responses. From a continual learning perspective, there has been recent work which identified algorithmic insights in the feedforward path of this system [12]. The full connectivity of the mushroom body is complex though, with specific recurrent connections of protocerebral anterior medial (PAM) neurons in the *α*1 compartment having been identified to be necessary for long-term memory (LTM) formation [5]. It is unclear, however, how these PAM-*α*1 feedback connections are enabling memory consolidation from an algorithmic perspective.

Given the benefits of replay-based approaches to continual learning in artificial systems and the theorized role of generative replay in higher level mammalian systems [3], our work here is motivated by two questions: 1) Is generative replay utilized for continual learning in this recurrent circuit for long-term memory formation? 2) If so, can this circuit yield new insights into principles for how to structure artificial systems for continual learning? In this work, we first investigate synapse-level connectivity data for evidence that generative replay may be the algorithmic mechanism supporting the consolidation of long-term memory. Next, we introduce a computational model for how generative replay may be utilized in this circuit to support continual learning. Finally, we evaluate the model’s performance in a CIFAR-100 [6] class-incremental baseline learning task through a series of experiments to obtain a more detailed understanding of the system’s properties.

## 2 Analysis of Connectivity for Evidence of Generative Replay Mechanism

We analyze the synapse-level connectivity released for an adult *Drosophila melanogaster* fruit fly [11] (which includes the full right mushroom body) for insights into whether or not generative replay may be the mechanism underlying long-term memory formation in the α1 compartment. To start, we focus our analysis on MBON07 and PAM11 which are the MBON and PAM types that have their primary arborization in the α1 compartment. If generative replay is utilized, we would expect that the patterns of sensory encoding neurons (KC’s) replayed for a certain memory would be similar to the patterns in the initial experience that are linked to the behavioral response. In this circuit, the PAM-KC connectivity can support reactivating the KC’s and the KC-MBON can support learning a behavioral response from KC activation. The patterns of connectivity at KC-MBON and PAM-KC synapses are measured as the count of synapses as a proxy for the effective coupling strength or weight between neurons as *W_KC→MBON_* and *W_PAM→KC_*. This connectivity can be compared along the KC dimension by measuring the pairwise correlation for each of the two columns (for two α1 MBON’s) in *W_KC→MBON_* and the seven rows (for seven α1 PAM neurons) in 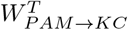 as well as the correlation between the average of the columns and rows respectively. When compared (Fig. 1), we find that *W_KC→MBON_* and 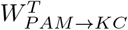 are correlated (relative to a null shuffled hypotheses) and therefore support the theory that the PAM-KC connections could support activating similar patterns as those encoding the initial sensory experience. This insight can also be interpreted as a property of the weight values of this circuit, which we incorporate into our replay model. Note that there is a relatively direct mapping between MBON07 and PAM11 connectivity, where over 85% of the MBON07 outgoing synapses to other PAM neurons go to a small population of PAM11 neurons and thus the MBON-PAM connectivity can be presumed to have a smaller role in shaping feedback activation than PAM-KC connectivity in the MBON-PAM-KC loop.

**Figure 1:**
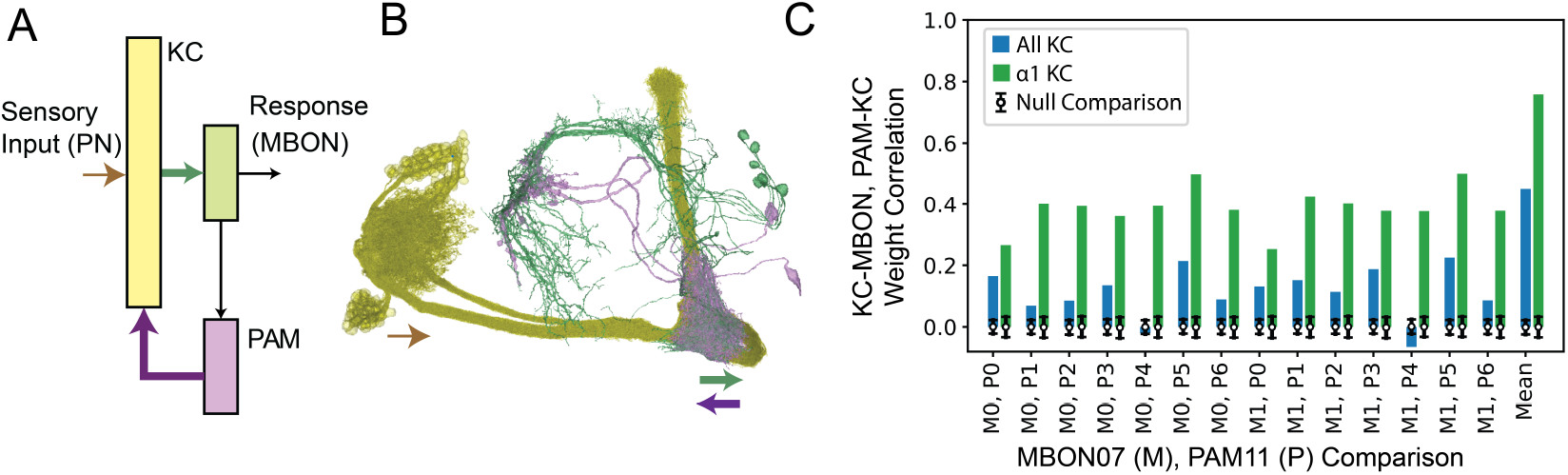
Connectivity analysis for evidence of generative replay. A. Neuron populations involved in a recurrent circuit found to drive appetitive LTM formation between a sensory input pattern and learned response in the α1 compartment of the mushroom body. The KC-MBON and PAM-KC connections are necessary for LTM formation and are theorized here to support continual learning via generative replay. B. Synapse-level connectivity data is available for this circuit [11] enabling interrogation of the circuit mechanisms enabling LTM. The sites of synaptic connections are annotated below the skeletons of the neurons. C. The count of synapses between neurons are used as a proxy for weight between neurons and analyzed for the KC-MBON and PAM-KC connections (*W_KC→MBON_*, *W_PAM→KC_*). The correlation between *W_κC→MBON_* and 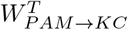 is measured for each pair-wise comparison between neurons (and the average across neurons) utilizing either all KC’s or α1 compartment KC’s. The correlation is found to be significant compared to a null condition with shuffled KC indices for 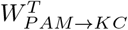 (error bars are standard deviation from 1000 random shuffled null comparisons with permuting the KC indices of *W_PAM→KC_*).

## 3 Generative Replay Model

We introduce an artificial neural network model that captures the connectivity in this circuit and investigate generative replay as a computational mechanism for long-term memory formation (see Fig. 2). Activation of the sensory-encoding KC’s (*h* ∈ ℝ^*D_h_*^) at timestep (*t*) are modeled as

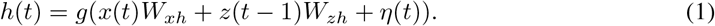

**Figure 2:**
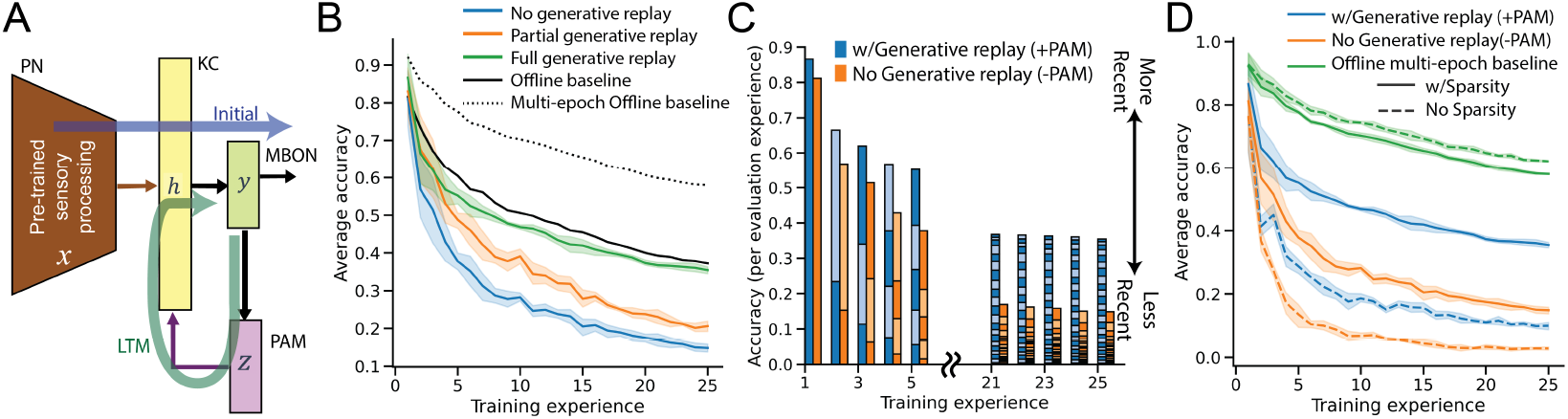
Continual learning model and experimental results. A. Initial experience is modeled by training the feedforward portion of this circuit given processed sensory input and a teaching signal, Long-term memory (LTM) with generative replay is modeled by training the recurrent portion of this circuit to reconstruct a sample from the output of the network. B. Average accuracy in a CIFAR-100 class-incremental learning task. C. Average task accuracy composition across training experiences (4 learned classes per experience). The number of stacked bars for each training experience is the number of evaluated previous experiences. Adding generative replay has less of a recency bias. D. Average accuracy with and without k-winners-take-all sparsity. Sparsity has greatest increase on generative replay accuracy (and reduces offline baseline accuracy). Error bars are bootstrapped 95% confidence interval of 5 random initialization seeds.

Network input to h includes processed sensory input (*x* ∈ ℝ^*D_x_*^) and PAM feedback activation (*z* ∈ ℝ*^D_z_^*), with associated network weights *W_xh_* and *W_zh_* respectively. A noise term 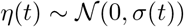 is also included in the model to introduce stochasticity. A non-linearity, *g*(.) enforces sparse activation using k-winners-take-all. The MBON activation, *y*(*t*) = *h*(*t*) *W_hy_*, is the model output based on network weights, from KC activation. Finally, PAM activation is calculated based on MBON activation, *z*(*t*) = *y*(*t*)*W_yz_*. Further insight from the synapse count correlation analysis in Fig. 1 can be captured through constraining 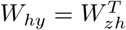. The network can be applied to a sensory pattern classification task, by adapting the dimensionality of *y* to the number of target classes.

To model the learning process during initial experience as a feedforward system (as has been investigated previously [1; 8; 12]), we utilize a single timestep with input *x*(0) and a teaching signal 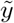 without PAM feedback or a noise term, *h_init_* (0) = *g*(*x*(0) *W_xh_*). The process of learning during initial experience is thus the update of model weights *W_hy_*, which can be performed with associative learning rules or by explicitly minimizing an objective function loss (*L_init_*) of *C*, 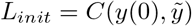.

The learning process in the long-term memory phase is modeled with the absence of sensory input, *h_LTM_*(*t*) = *g*(*z*(*t* − 1)*W_zh_* + *η*(*t*)). The generative replay in this system is driven by a sampled MBON activation pattern, *y_s_*(*θ*), which is all zero except for a randomly chosen single unit (one-hot embedding). Learning in this long-term memory phase is performed by updating network weights *W_hy_* and *W_zh_*, which can be performed again via associative learning rules or by explicitly minimizing an objective function loss (*L_LTM_*) of *C*, *L_LTM_* = *C*(*y_s_*(0), *y*(1)).

## 4 Experiments

To investigate performance of the generative replay model, we extend the modeling and continual learning investigation approach utilized for the studying the feedforward portion of this model [12]. A class-incremental learning CIFAR-100 baseline task (25 training experiences of 4 classes each) is used where processed sensory input is Imagenet pre-trained Resnet embeddings (dimensionality 512), as further detailed in [12]). A 10% sparse binary matrix is used for *W_xh_* and the nonlinearity *g*(.) enforces k-winners-take-all, with k equal to 1% of *D_h_*, where *h* has a 40 × expansion in dimensionality relative to *x*. To isolate our investigation on the generative replay portion our model, we explicitly minimize the objective function loss terms, *L_init_* and *L_LTM_* utilizing stochastic gradient descent of the cross entropy loss (learning rate=1e-3, momentum=0.9, batch size=32 and 64 for initial experience and LTM respectively). Investigation of associative learning rules based on coactive neurons for updating network weights is left to future work. To investigate the implications of the connectivity trends in the anatomical data, where *W_hy_* is correlated with 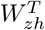, the weight values are tied during optimization to 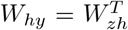. Additional simplification includes modeling the *W_yz_* MBON-PAM connectivity as identity based on the observed relatively direct mapping between MBON and PAM connectivity as further described in the previous section. All code implementing additional model and training details will be released at https://github.com/aplbrain/mb_gen_replay.

We measure the relative effect of the proposed LTM by comparing average accuracy. An upper bound baseline on CIFAR-100 (without class-incremental learning) and several repeats of input data (50 epochs) is reduced with only one presentation of the data (1 epoch), (average final accuracy from 0.58 to 0.37). This performance is drastically reduced when introducing the class-incremental presentation of training data (from 0.37 to 0.15). When the long-term memory phase of the model is utilized after each training experience, there is a marked recovery in accuracy (from 0.15 to 0.35) which approaches the non-incremental training of the task (average final accuracy of 0.37).

To further investigate the properties of the modeled MB network LTM learning phase, we investigate the balance of accuracy between experiences as a function of recency. Adding the LTM training phase, the accuracy is markedly more balanced between experiences, with less bias towards performance on more recent experiences (Fig. 3A). A property of the MB network is that there are sparse activation in KC’s. When this modeled sparsity is removed, there is actually an increase in the offline (non-class incremental) version of this task. Sparsity is instrumental though in enabling performance in the generative replay model (jump of final accuracy from 0.10 to 0.35).

## 5 Discussion

In this work, we propose a theory that the algorithmic mechanism behind LTM formation in the α1 compartment of the mushroom body is generative replay and identify evidence of this plausibility in synapse-level connectivity data. We introduce a novel computational model of mushroom body LTM formation using generative replay and find the addition can increase performance on a class-incremental learning CIFAR-100 task to the level of an offline upper bound baseline. Furthermore, the performance is competitive with other biologically-inspired approaches for continual learning [12; 13] and could potentially be used in future machine learning systems.

To inform development of future continual learning systems, there are a few particularly distinctive properties of this system that could be leveraged more broadly. For one, the generative replay mechanism is not optimized based on storing processed sensory input [2] or learning to generate processed sensory input [13], which incur memory costs for storing data from past tasks or the current task (respectively). Instead, optimization of the generative replay component is performed based on sampling a higher-level representation and optimized to reconstruct the higher-level representation created in a recurrent feedback loop in the system (Fig. 2A). An additional component utilized in this system which was shown to be necessary for performance is a high-dimensional, sparsely activated intermediate encoding layer. Interestingly, the sparse activation caused a decrease in offline comparison, but was instrumental in continual learning, thus emphasizing that network properties beneficial for traditional multi-epoch offline training may be detrimental to performance in continual learning.

The evaluated synapse-level connectivity offers intriguing evidence that generative replay may be an algorithmic mechanism utilized for LTM consolidation in the fruit fly, but is far from conclusive. A major contribution of this computational study is the experimentally testable prediction that the PAM to KC synapses in the α1 compartment undergo experience-dependent plasticity and that this experience-dependent plasticity is necessary for LTM formation.

Future work could further investigate the computational principles underlying LTM formation in this circuit, utilize components of this model in more complex artificial systems, or test this hypothesis experimentally in the fruit fly.

## Acknowledgments and Disclosure of Funding

We thank Yang Shen and Saket Navlakha for their helpful discussion and for sharing details of their fly continual learning model. This research received internal funding from the Johns Hopkins University Applied Physics Laboratory.

